# A Comprehensive Atlas and Machine-Learning Framework for Predicting IDR-Protein Binding Affinity

**DOI:** 10.64898/2026.02.21.707165

**Authors:** Subinoy Adhikari, Soham Choudhuri, Jagannath Mondal

## Abstract

Intrinsically disordered regions (IDRs) enable pervasive, regulatable protein interactions, yet predicting their binding affinities remains challenging because disorder permits heterogeneous interfaces and context-dependent recognition. Here we introduce **IBPC-***K_d_*, a curated dataset of 1,785 IDR-ordered protein complexes with experimentally measured dissociation constants (*K_d_*), spanning more than six orders of magnitude in affinity and covering diverse IDR lengths and partner classes. Across this resource, we find that global interfacial shape complementarity is a consistent correlate of affinity, followed by the structural order of the partner surface and a systematic electrostatic asymmetry between IDRs and their binders. Local physicochemical patterning modulates binding in a partner-dependent manner. Guided by these observations, we develop **IDRBindNet**, a graph-transformer model that integrates protein language model embeddings with interface geometry and residue-level chemical context to predict dissociation constants from IDR-partner complexes. **IDRBindNet** achieves state-of-the-art accuracy on held-out complexes (R^2^ up to 0.911) while implicitly learning biologically relevant attention patterns corresponding to partner disorder and geometric fit. Notably, the framework generalizes through external validation on a set of recently reported *de novo* designed binders targeting the human surfaceome, showing encouraging concordance across nanomolar-to-micromolar affinities. Together, **IBPC-***K_d_*and the accompanying predictor **IDRBindNet** provide a quantitative framework for interrogating determinants of IDR recognition and for prioritizing candidate interactions for mechanistic and design studies.

## Introduction

Intrinsically disordered proteins and intrinsically disordered regions (IDRs) play essential roles in biological systems, particularly in regulatory processes, signaling networks, and dynamic protein-protein interactions.^1^ A major class of IDRs comprises the activation domains and regulatory regions of transcription factors (TFs), where structural plasticity enables interactions with co-activators, co-repressors, mediator complexes, and other partners to control gene expression.^1,2^ Unlike globular proteins, IDRs lack a stable tertiary structure in their isolated form, yet they frequently adopt ordered conformations upon binding to partner proteins.^3^ These disorder-to-order transitions enable distinct interaction modes that differ from classical globular protein complexes, often involving short, flexible segments that form specific structural elements in the bound state.^4^

Classic examples include the binding of the intrinsically disordered transactivation domain of p53 to the globular protein MDM2, where the IDR folds into an amphipathic alphahelix upon interaction; ^5^ the disordered region of p27 (Kip1) binding to the cyclin A-CDK2 complex, adopting a defined structure in the bound state to inhibit kinase activity;^6^ the interaction of E-cadherin’s disordered cytoplasmic tail with beta-catenin, involving coupled folding and binding;^7^ and the disordered regulatory domain of eIF4G engaging the globular eIF4E to regulate translation initiation.^8^ These cases illustrate how IDRs exploit structural plasticity to achieve high specificity despite initial flexibility, often resulting in interfaces that differ from classical globular-globular complexes by relying on induced folding and extended contact surfaces. ^2,9^

The structural plasticity and lack of fixed conformation in IDRs have historically made them challenging targets for conventional drug discovery approaches, often rendering them “undruggable”. However, recent advances, particularly from computational protein design strategies developed in Baker’s^10,11^ lab have opened new possibilities for therapeutic intervention. Using tools like RFdiffusion^12^ and modular assembly methods^10^ (e.g., “logos”), high-affinity binders have been de novo designed to target diverse IDRs, achieving nanomolar dissociation constants^10,11^ and demonstrating applications in disrupting pathological interactions, dissolving amyloid aggregates, or modulating signaling pathways. These designed binders highlight the potential to selectively target IDR-mediated complexes in transcription factors and other regulatory proteins, paving the way for novel diagnostics and therapeutics against diseases involving dysregulated transcription, signaling, or aggregation-prone IDRs.

Structural studies of selected IDR complexes have revealed unique binding principles and facilitated the development of specialized prediction tools for disordered binding sites. ^13,14^ Complementary approaches, such as the identification of short linear motifs (SLiMs), have further characterized functional binding determinants within disordered regions. ^15^ Despite these advances and occasional success in pharmaceutical targeting of IDR interactions^16^ systematic, high-resolution understanding of the biophysical principles governing IDR binding remains limited. Existing databases have provided valuable resources but address only specific aspects of this landscape. Sequence-centric repositories such as DisProt, ^17^ MobiDB^18^ and IDEAL^19^ catalog disorder at the primary structure level, while interaction-focused resources like DisBind,^20^ ELM,^21^ FuzDB^22^ and MFIB^23^ and structure-centric resources like PDBtot and PDBcdr^24^ either prioritize sequence motifs or focus on IDR-IDR complexes. However, these resources generally lack comprehensive quantitative data on binding affinities, which are critical for linking structural features to thermodynamic and functional outcomes, for benchmarking next-generation prediction algorithms, and for exploring relationships between protein disorder and disease.

To bridge this gap, we present **IBPC-***K_d_*, a large-scale dataset of IDR-Binder Protein Complexes interactions with experimentally determined dissociation constants (*K_d_*). At the time of this study, the DIBS database represented the only publicly available resource that systematically reports *K_d_* values for IDR-protein interactions. However, DIBS^25^ contains 772 entries out of which only 487 entries reported *k_d_* values which were compiled from earlier experimental studies, making its size insufficient for training machine learning models without a substantial risk of overfitting. Starting from these 487 DIBS entries, we expanded the collection through manual curation of high-quality affinity measurements from diverse published studies. After careful deduplication (DIBS data was kept as such), the final dataset comprises 1,785 unique IDR-protein pairs with associated *K_d_* values. Building on this dataset, we developed **IDRBindNet**, to our knowledge the first predictive model for estimating *K_d_* of IDR-ordered protein interactions. Our approach uses a hybrid graph-based representation of each complex, integrating embeddings from protein language models with structural features. A transformer convolutional architecture then processes this representation, employing an attention mechanism to learn the determinants of binding. Notably, the model’s attention heads provide intrinsic interpretability, enabling us to identify which structural and sequence features underlies strong and weak binding affinities.

## Results

### IBPC-*K_d_*: Curated dataset of IDR-protein binding affinity

We started curating an expanded dataset of IDR-protein interactions with experimentally determined *K_d_*values, starting from the DIBS database^25^ as a foundational resource. Building upon this legacy curation, we systematically integrated additional data from multiple specialized sources to broaden both the affinity range and biological diversity of the dataset. These include high-affinity binders generated by deep-learning-based protein design from the Baker group,^10,11^ quantitative analyses of well-defined biological systems such as the calcineurin-phosphopeptide^26^ and DREB2A-RCD1 complexes,^27,28^ systematic affinity measurements from high-throughput combinatorial peptide libraries (the N2P2^29^ dataset), and large-scale profiling of human transcription factor activation domains against multiple coactivators (the STAMMPPING dataset^30^). As shown in Figure 1 (a), the composition of our curated dataset, referred as **IBPC-***K_d_*, is illustrated in a pie chart highlighting the proportional contribution of each source. The aggregated data reveals several defining characteristics: the experimentally measured affinities span more than six orders of magnitude, from approximately 1 nM to over 100 *µ*M (see Figure1 (b)), capturing the full physiological spectrum of interaction strengths. The binder proteins are functionally diverse, encompassing enzymes, scaffolds, and components of the transcriptional machinery to ensure broad biological relevance. The IDRs themselves vary from short linear motifs of fewer than 10 residues to longer disordered domains of approximately 80 residues, enabling a systematic investigation of affinity determinants across multiple length scales of disorder.

**Figure 1:**
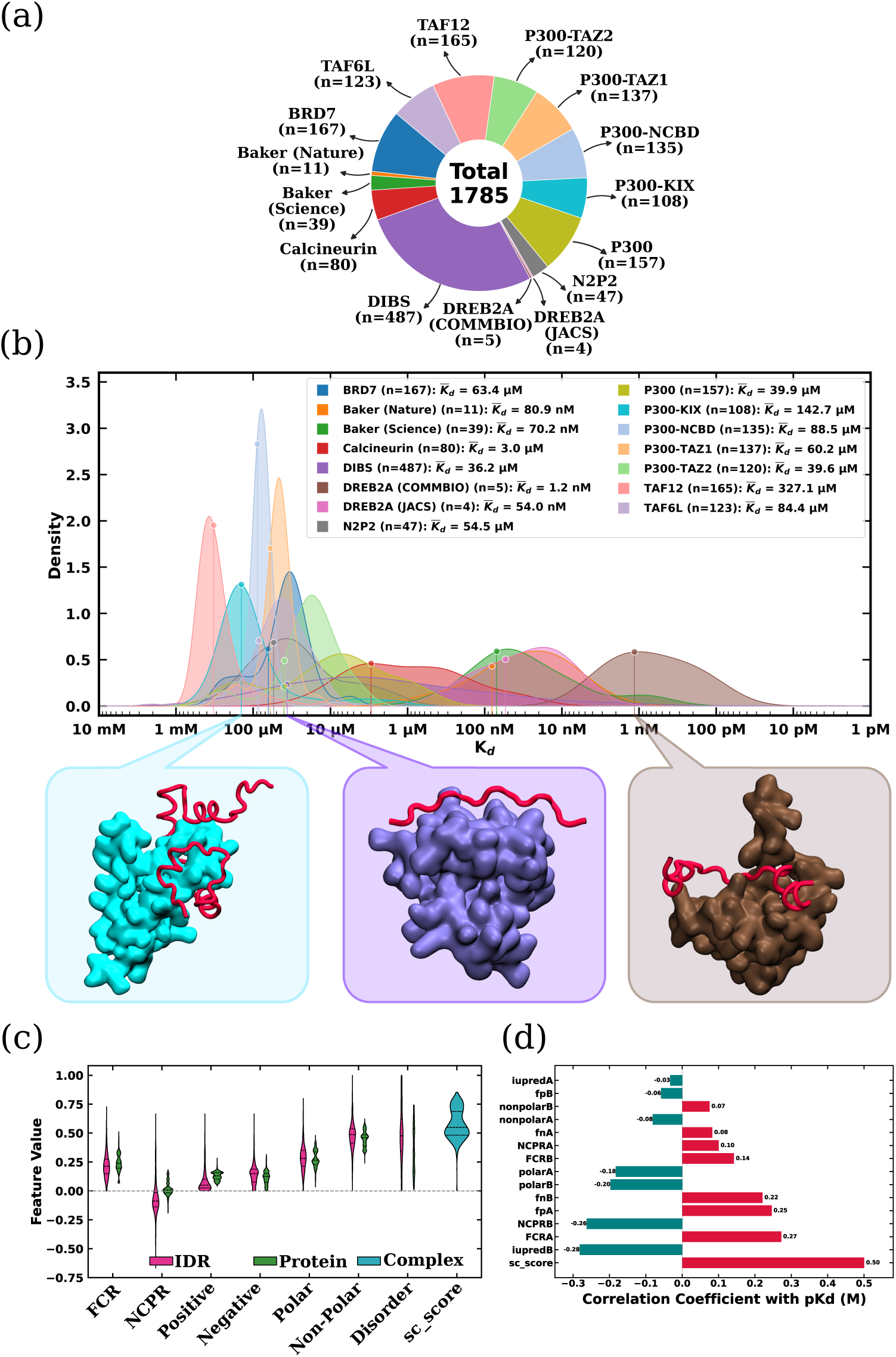
*IBPC* − *K_d_*data-set in a nutshell: (a) Pie chart showing proportional composition of various sources that constituted *IBPC* − *K_d_* (b) Distribution of *K_d_* values across different datasets along with the representative structures present in *IBPC* − *K_d_* (c) Distribution of physicochemical and strctural properties (d) Correlation of *pK_d_*with physicochemical and strctural properties in *IBPC* − *K_d_*datasets.

This expansive range of affinities as presented by **IBPC-***K_d_* is reflected in the distribution of *K_d_*values (see Figure1 (b)). The DIBS dataset exhibits one of the broadest and most heterogeneous affinity distributions, characterized by a strong mean-median separation. Its mean *K_d_* is in the tens of micromolar range (∼36 *µ*M), while its median is an order of magnitude lower (∼3 *µ*M), with a large standard deviation (∼152 *µ*M). This indicates a highly skewed distribution dominated by many weak interactions, alongside a smaller subset of tighter binders. In contrast, other datasets occupy distinct affinity regimes. At the strongest end, the DREB2A^27,28^ datasets shows uniformly tight binding, with a mean *K_d_* of ∼1 nM and a very small standard deviation. The DREB2A and Baker^10,11^ datasets also fall within the high-affinity nanomolar regime (tens of nanomolar) but with noticeably larger standard deviations relative to their means, reflecting a mix of strong designed binders and weaker variants from mutational exploration. The calcineurin data^26^ resides in an intermediate regime, with a mean *K_d_*around ∼3 *µ*M. The BRD7,^30^ N2P2,^29^ and P300^30^ datasets cluster in the mid-to-weak affinity range, with means in the tens of micromolar and with large standard deviations. The weakest average affinities are observed for the P300-KIX ^30^ and TAF12^30^ datasets, with means between ∼150-300 *µ*M and relatively narrower spreads. Finally, the P300-NCBD, ^30^ P300-TAZ1,^30^ P300-TAZ2, ^30^ and TAF6L^30^ datasets occupy a moderate micromolar regime, where medians are often lower than means, indicating similarly skewed distributions with subsets of tighter binders(see Figure 1 (b)).

### Global physicochemical determinants of IDR-protein binding

Each IDR-binder complex in *IBPC*−*K_d_* data-set was represented using a set of physicochemical and structural descriptors designed to capture the key biophysical principles governing molecular recognition by IDRs. ^2,31^ These descriptors were chosen based on their established relevance in characterizing the sequence properties, binding-coupled folding, and interfacial features that are characteristic of IDR-mediated interactions; descriptors with the suffix A correspond to properties of the IDR, while those with the suffix B refer to the binder protein.

Electrostatics plays a central role in determining both affinity and specificity in IDR binding.^32–34^ To quantify these charge-related effects, we included the fraction of positively charged residues (fpA, fpB), negatively charged residues (fnA, fnB), the overall fraction of charged residues (FCRA, FCRB), and the net charge per residue (NCPRA, NCPRB). Collectively, these metrics capture the electrostatic character of IDR and the binding partner, which is often a primary driving force for complex formation.^35,36^

The degree of intrinsic disorder in both the IDR and its binding partner was quantified using averaged IUPred2A scores^37^ (iupredA, iupredB). These energy-based scores estimate the propensity of residues to remain disordered in isolation and serve as a reliable proxy for intrinsic folding tendencies, thereby providing insight into the likelihood and extent of binding-coupled folding upon complex formation. To characterize the chemical nature of the interaction interface, we computed the fractions of polar (polarA, polarB) and nonpolar (nonpolarA, nonpolarB) residues. These descriptors distinguish between interfaces dominated by hydrophilic, solvation-driven interactions and those involving hydrophobic burial, which are associated with distinct thermodynamic signatures.^38–40^ Finally, the geometric complementarity of the complex was captured using the shape complementarity score (sc_score) from PyRosetta. ^41^ This structural descriptor enables the correlation of binding affinity with the three-dimensional steric fit achieved upon binding, a well-recognized contributor to complex stability. ^42^ Together, this fifteen dimensional feature set integrates sequence-based, biophysical, and structural information to provide a framework for analyzing the determinants of IDR-protein interaction affinity.

To understand how these features collectively shape the affinity landscape of IDR-mediated interactions, we analyzed the distributions of the physicochemical descriptors across the *IBPC* − *K_d_* dataset (see Figure 1 (c)). Charge-related features reveal complementary trends between the two chains. Although the FCR is comparable, the binder protein shows a slightly higher median FCR (0.24) than the IDR (0.21), indicating that binders are also charge-rich but in a more balanced manner. This distinction becomes clearer when examining the NCPR values: the IDR displays a predominantly negative distribution with a median NCPR of -0.09, whereas the binder protein is centered near neutrality with a slightly positive median (0.008). Decomposition into positive and negative charge fractions further highlights this asymmetry, as the IDR is enriched in negatively charged residues (median 0.15) and depleted in positively charged residues (median 0.05), while the binder protein shows the opposite trend, with a markedly higher fraction of positively charged residues (median 0.13) and a lower negative fraction (median 0.12). These distributions indicate that electrostatic complementarity between negatively biased IDRs and positively biased folded partners is a dominant feature of the interaction landscape. Analysis of residue composition shows that both chains are dominated by non-polar residues, but with subtle differences. The IDR exhibits a slightly higher median non-polar fraction (0.49) compared to the binder protein (0.46), while also maintaining a marginally higher polar fraction (0.28 vs. 0.27). Disorder scores derived from IUPred2A show a clear separation between the two chains, with the IDR exhibiting a substantially higher median disorder (0.48) than the binder protein (0.35), confirming the intrinsically disordered nature of the IDR and the comparatively ordered character of the binder proteins. Finally, shape complementarity, for the IDR-protein complex, shows a distribution with a median value of 0.55 (see Figure 1 (c)).

Correlation of these features with *pK_d_* (*pK_d_* = −log_10_*K_d_*) reveals a clear hierarchy of determinants. The strongest correlation is observed for shape complementarity, indicating that the degree of geometric matching at the interface is the primary global determinant of binding strength. The second strongest association is a negative correlation with disorder in the binder protein, suggesting that increased flexibility in the folded partner weakens binding. The fraction of charged residues in the IDR, along with the fraction of positively charged residues in the IDR and negatively charged residues in the binder protein, shows moderate positive correlations with affinity, highlighting a supportive role for electrostatics. Collectively, these results establish shape complementarity, structural order of the binder, and charge asymmetry as the dominant global features governing IDR-protein binding (Figure 1(d)).

A particularly striking contrast emerges when comparing complexes originally obtained from the Baker and DIBS sources, which occupy markedly different affinity regimes. While the Baker source is dominated by high-affinity interactions in the nanomolar range, the DIBS source is centered in the micromolar regime. To understand the origin of this disparity, we examined dataset-specific correlations between *pK_d_* and the physicochemical features described above. Notably, the Pearson correlation coefficient (PCC) for shape complementarity and for the fraction of non-polar residues in both the IDR and the binder protein are substantially higher in the Baker dataset than in DIBS, indicating a stronger coupling between hydrophobic packing, geometric matching, and affinity in the rationally designed systems. (see Figure S1)

### Latent interaction regimes revealed by unsupervised clustering

While the feature-wise analysis identifies global determinants of binding affinity, it does not address whether IDR-protein interactions segregate into distinct physicochemical regimes characterized by different combinations of these features. To address this, the 15-dimensional feature space was reduced to two dimensions using Principal Component Analysis (PCA) (see Figure S2), t-distributed Stochastic Neighbor Embedding (t-SNE)^43^ (see Figure S2), and Uniform Manifold Approximation and Projection (UMAP)^44^ (see Figure 2 (a)), of which UMAP yielded the clearest separation (see “Methods” for further details). The resulting clusters occupy distinct and well-separated regions of the UMAP space, with clearly defined cluster centers (Figure 2 (c)). Importantly, these clusters correspond to distinct binding affinity regimes. The mean *pK_d_* varies substantially across clusters, spanning approximately 3.67 to 6.00, indicating that the unsupervised partitioning captures meaningful differences in interaction strength. To interpret the physicochemical basis of these regimes, we computed cluster-wise z-scores for all features (Figure 2 (d)). Each cluster exhibits a distinct feature signature: some are strongly enriched in shape complementarity (notably cluster 8), others in charge content of the IDR (clusters 0, 1, and 5), polarity differences (clusters 4 and 7), or varying degrees of disorder in either the IDR (cluster 6) or the binder protein (clusters 2 and 3). To connect these regime-level signatures to binding affinity, we correlated cluster-wise feature z-scores with the corresponding mean *pK_d_* values. Shape complementarity emerges as the dominant determinant at the cluster level (*r* = 0.90), indicating that clusters enriched in well-packed interfaces consistently exhibit stronger binding. Charge-related features of the IDR show moderate positive correlations, suggesting that electrostatics modulate affinity within specific interaction regimes rather than acting as a universal determinant. In contrast, disorder in the binder protein is negatively correlated with *pK_d_*, reinforcing the importance of a structured binding surface. Notably, several compositional features of the binder protein exhibit weak or negligible correlations at the cluster level, underscoring that geometric complementarity, rather than residue composition alone, governs high-affinity IDR-protein interactions across distinct interaction modes (see Figure S3).

**Figure 2:**
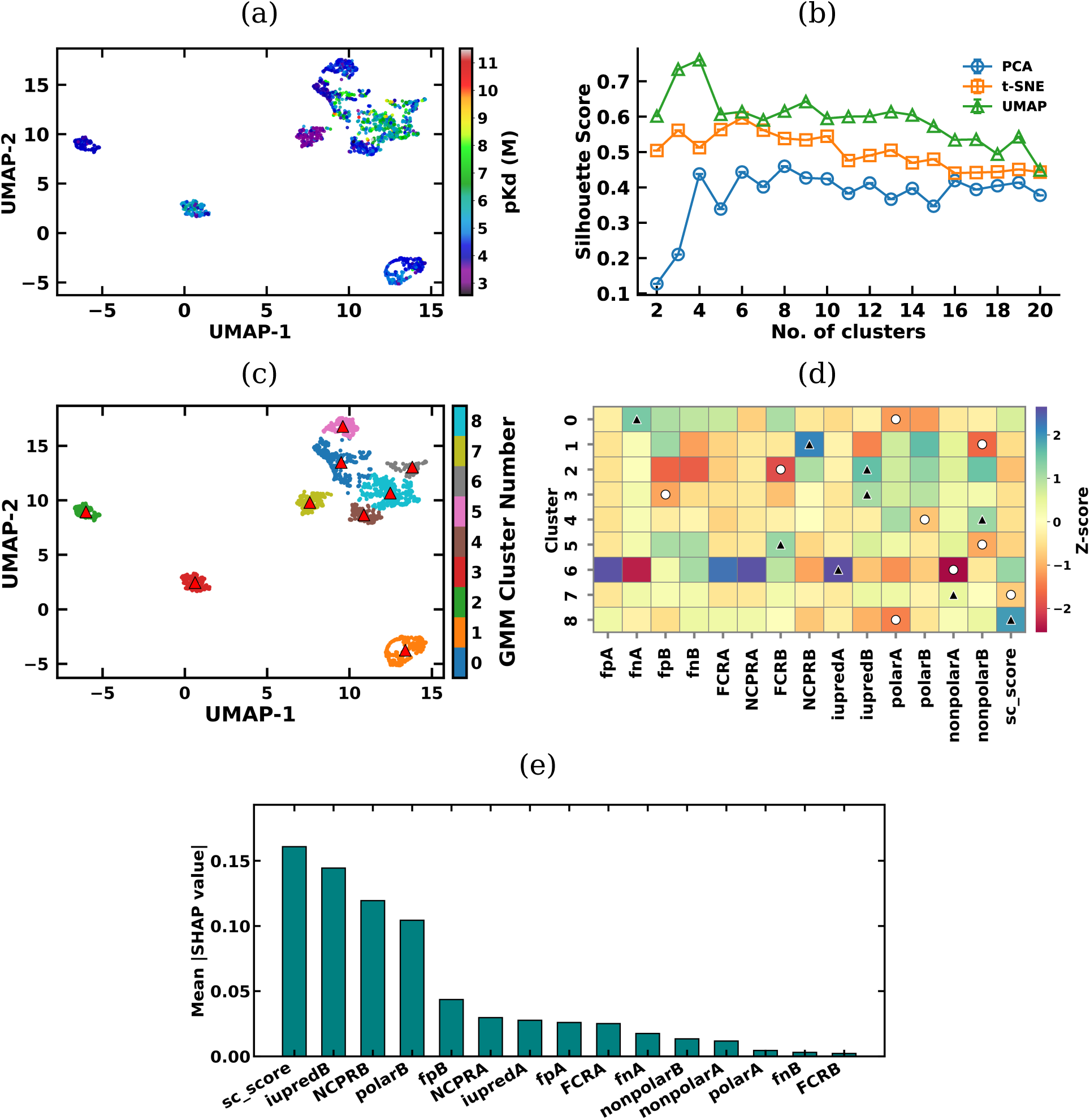
(a) Latent structure of the dataset visualized using UMAP. (b) Comparison of silhouette scores across different numbers of clusters for PCA, t-SNE, and UMAP embeddings. (c) Cluster assignments obtained using GMM clustering. (d) Z-score-normalized physicochemical and structural features across the identified clusters. White circles and dark trainles correspond to minimum and maximum z-scores. (e) SHAP analysis of the Random Forest regression model for *pK_d_* prediction.

While clustering of *IBPC* − *K_d_* data-set reveals groups of complexes with different binding affinities, it does not directly assess how well these features can quantitatively predict *pK_d_* for individual complexes. To address this, we next trained supervised regression models using the same features to explicitly predict binding affinity. Linear regression models (Linear, Lasso,^45^ and Ridge^46^) showed only modest performance on the test set, with *R*^2^ values around 0.24-0.25. The fact that all three gave almost the same result even after tuning regularization for Lasso and Ridge suggests that *pK_d_* cannot be well explained by simple linear combinations of the features we used. In contrast, the nonlinear ensemble models performed much better. Both XGBoost^47^ and Random Forest^48^ regression models reached *R*^2^ values of about 0.68-0.69 which is almost three times higher than the linear models (see Figure S4). This large improvement shows that *pK_d_* depends strongly on nonlinear interactions between features. To understand which features mattered most, we applied SHAP^49^ analysis to the best-performing Random Forest model. Shape complementarity was by far the most important feature, showing the highest mean absolute SHAP value. This confirms that how well the shapes of the two proteins fit together is the main driver of binding strength. Disorder in the binder protein and the binder’s net charge per residue were also among the top features. Together, these results indicate that a structurally ordered binding surface with favorable electrostatics is important for strong binding.

Nevertheless, the achieved *R*^2^ values are still low to provide a reliable prediction of binding affinities. Furthermore, these models rely on a fixed set of global, hand-crafted descriptors and therefore cannot fully capture the complex residue-level interactions that underlie IDR-protein binding. Recent advances in protein language models provide rich, sequence-informed representations that encode both local and long-range dependencies, offering an opportunity to move beyond coarse-grained features. Motivated by this, we developed a graph-based transformer model, coined as **IDRBindNet**, that integrates protein language model embeddings with explicit structural information to improve the prediction of binding affinity.

### Predicting binding affinity with IDRBindNet

The architecture of **IDRBindNet** is illustrated in Figure 3. Graph-based models are naturally well suited for learning from datasets containing IDR-protein complexes of varying sizes, as they operate on relational structure rather than fixed-length representations. This allows complexes of different lengths represented by *IBPC* − *K_d_* datasets to be processed without the need for truncation or padding. At its core, the model processes a residue-level graph representation of the complex where each node corresponds to an amino acid residue. The node features are derived from protein language models (PLM), for which we explored multiple variants of PLMs including Evolutionary Scale Modeling (ESM-2)^50^ framework and the ProtTrans family,^51^ capturing evolutionary and functional constraints. The architecture leverages four key edge features to modulate this attention mechanism (see “Methods”): (i) the pairwise C*α*-C*α* distance establishes which residues are spatially proximate enough to interact; (ii) the relative orientation provides directional context for evaluating geometric complementarity; (iii) the difference in C*α* chemical shifts encodes local conformational and electronic environments that change upon binding; and (iv) the residue-wise difference in solvent-accessible surface area (SASA) reflects interfacial burial and hydrophobic driving forces (See details in “Methods”). By conditioning the attention mechanism on these descriptors, the model learns to prioritize interactions that are meaningful for predicting binding affinity.

**Figure 3:**
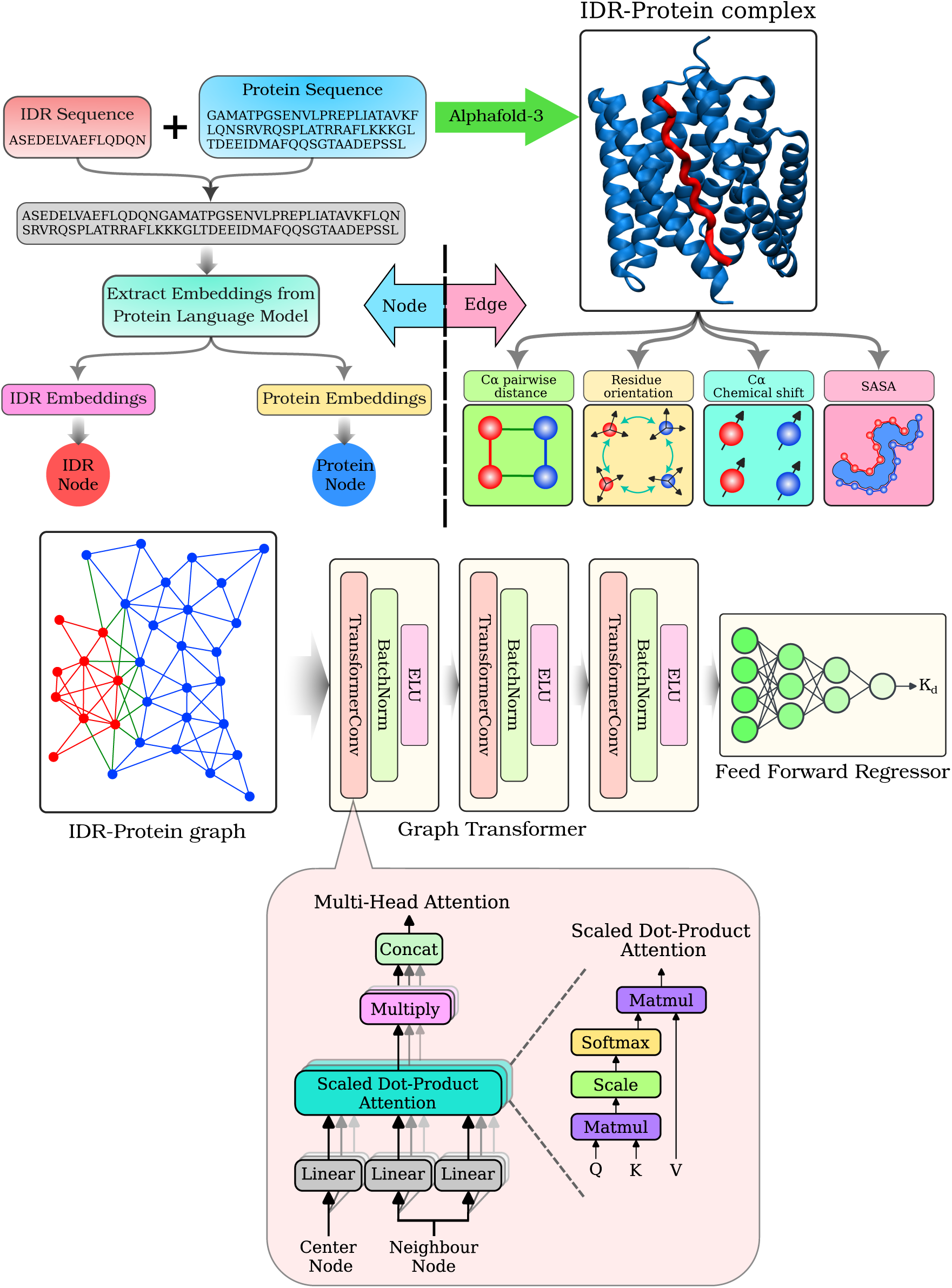
Model architecture of **IDRBindNet** to predict *K_d_*

We next evaluated the predictive performance of **IDRBindNet** to assess whether the architectural advantages described above translate into improved affinity prediction accuracy. Specifically, we compared sequence representations derived from different protein language models against established affinity prediction methods. Performance was assessed on an independent test set of 275 complexes using both the PCC and *R*^2^ values. Across all language-model embeddings, **IDRBindNet** achieved consistently high accuracy, with PCC values exceeding 0.92 and *R*^2^ values approaching or exceeding 0.90 for the largest models. Within the ESM2 family, performance improved systematically with increasing model size, with PCC increasing from 0.929 (ESM2-8M) to 0.947 (ESM2-3B) and corresponding *R*^2^ values reaching up to 0.895. The best overall performance was obtained using PROTT5-BFD embeddings (PCC = 0.956, *R*^2^ = 0.911), indicating excellent agreement between predicted and experimental *pK_d_* values. In contrast, existing generic protein-protein affinity predictors such as PPAP, despite achieving a high PCC, showed a substantially reduced *R*^2^ (0.582), while ProAffinity-GNN achieved moderate performance (*R*^2^ = 0.697), remaining significantly below our PROT-T5-BFD model (see Table S1). Consequently, **IDRBindNet** (see the Code Availability section) provides a first-of-its and efficient way to estimate the *K_d_* of an IDR-partner protein complex directly from its structure.

To establish a baseline and assess the contribution of learned sequence representations, we included a one-hot encoding of amino acid sequences as a control. Despite its simplicity and lack of contextual or evolutionary information, the one-hot model achieved strong performance (PCC = 0.924, *R*^2^ = 0.850), indicating that primary sequence composition alone carries substantial information relevant to binding affinity. However, all protein language model embeddings consistently outperformed the one-hot baseline, particularly in terms of *R*^2^, demonstrating that contextualized representations capture higher-order sequence patterns and long-range dependencies not accessible to naive encodings.

To further evaluate whether the observed performance reflects genuine generalization rather than residual sequence similarity bias, we implemented a stringent sequence-clustering split at a 40% identity threshold (see “Methods”). Conventional random train-test partitions can retain substantial homology overlap, potentially leading to inflated performance estimates driven by memorization of closely related sequences rather than true functional learning. By clustering sequences and ensuring that proteins sharing more than 40% identity were confined to the same fold, we enforced a rigorous sequence-level out-of-distribution (OOD) evaluation in which the model was tested on evolutionarily distant complexes. This strategy provides a more realistic assessment of predictive performance when applied to novel or uncharacterized protein families. Under these stringent conditions, **IDRBindNet** achieved a PCC of 0.879 ± 0.083 and an *R*^2^ of 0.757 ± 0.150 across folds. The relatively narrow standard deviation in PCC indicates consistent performance across diverse structural clusters, while the sustained *R*^2^ value under homology-controlled evaluation demonstrates that the architecture captures generalizable biophysical features rather than merely local sequence motifs.

Can our ML predictor retain accuracy when evaluated on complexes from a completely different source, outside the scope of our curated Protein-IDR dataset? Post-curation of our Protein-IDR affinity dataset for this manuscript, a new study by Balbi and coworkers^52^ reported a small but useful set of AI-designed, surface-bound protein complexes with experimentally measured *K_d_* values. This serendipitously offered a stringent external benchmark for our ML predictor, drawn from an entirely independent source and therefore free from any overlap with our curated data. Crucially, these are *de novo* designed protein binders engaging structured protein surfaces, i.e., a binding regime that was not present from the 1,785 IDR-protein complexes used to train our model, making this test an out-of-distribution challenge. Because these complexes were entirely absent from our training pipeline, they offered an ideal, unbiased benchmark. However, even when tested from a different source, our predictions via **IDRBindNet** track the experimental affinities with encouraging fidelity across multiple targets. In particular, the model show-cased encouraging prediction affinities from the micromolar range (2.5 *µ*M experimental vs. 7.3 *µ*M predicted) down to the nanomolar range (e.g., 172 nM experimental vs. 142 nM predicted) (see Table S3 and Figure S5). This successful blind validation underscores the robust predictive capabilities of our model and also highlights its exciting potential to well beyond the training dataset. Together, this independent benchmark supports that the model’s learned features are not narrowly tied to our curated Protein-IDR dataset, but remain predictive when confronted with unseen complexes, highlighting robustness under a genuine source and regime shift.

### Model attention aligns with shape complementarity and disorder

To gain deeper insight into the **IDRBindNet**’s internal behavior, we examined attention mechanisms across its two layers, each containing eight attention heads. Our analysis focused specifically on inter-chain edges corresponding to IDR-protein interactions. For each such edge, attention weights were collected and averaged across the entire dataset for every head. These attention profiles were then correlated with the set of 15 physicochemical and structural descriptors (see Figure 4(c)). In the first layer, we observed a distinct pattern: six of the eight attention heads showed strong and recurrent positive correlations with the intrinsic disorder of the binding partner, with peak values reaching up to 0.44. In contrast, features linked to electrostatic composition-particularly the fraction of positively charged residues in the partner sequence consistently exhibited the strongest negative correlations. In the second layer, most heads displayed their strongest negative correlations with the shape complementarity score, while retaining moderate positive associations with iupredB (see Figure 4 (d)).

**Figure 4:**
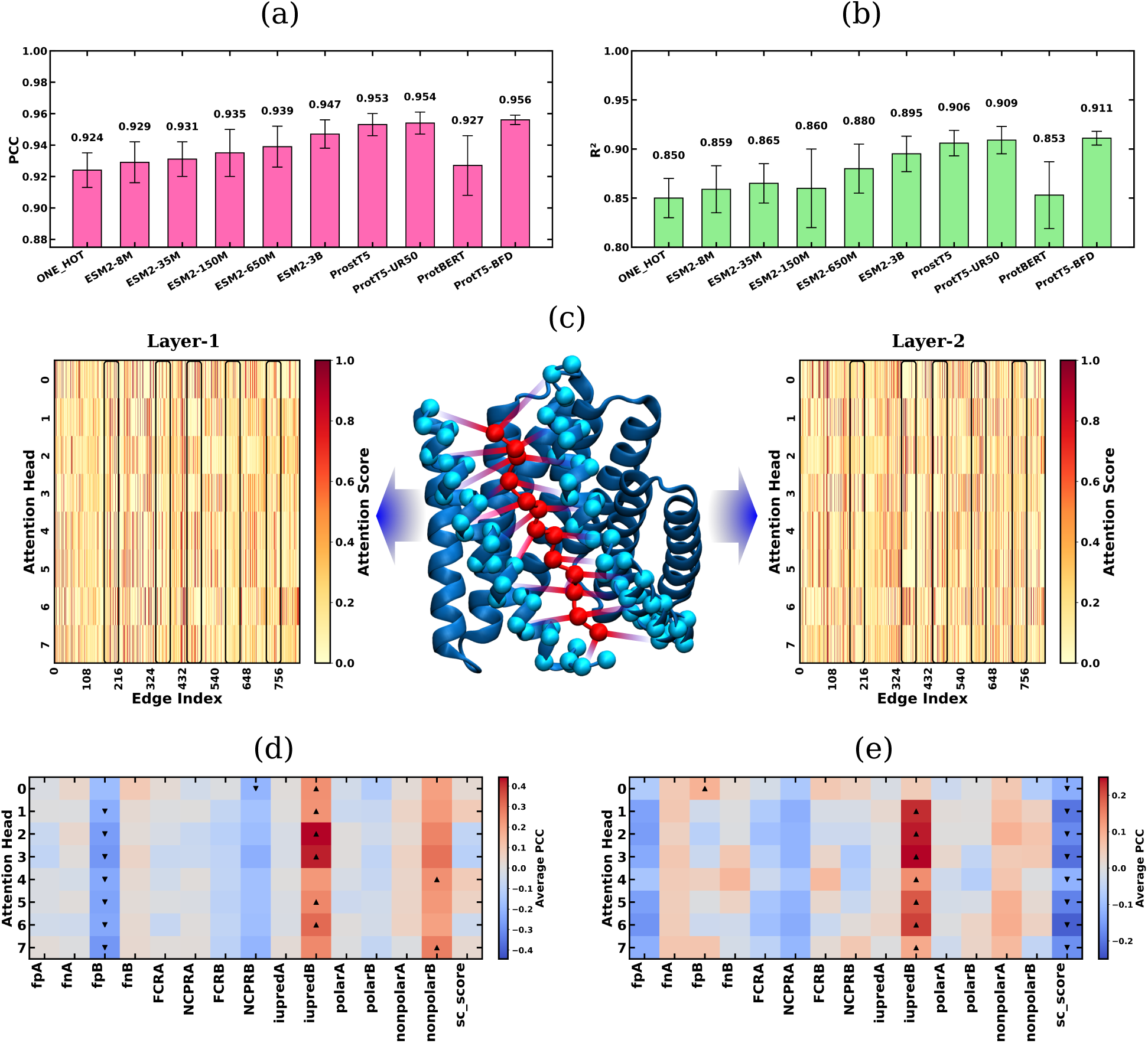
Comparison of (a) PCC and (b) *R*^2^ to predict *pK_d_* using different language models and baseline models tested in **IDRBindNet**. (c) Protocol for extracting average attention weights of edge features across attention head. Average attention weights across all heads in the (d) first layer and (e) second layer.

Notably, although none of the physicochemical or structural descriptors were used as input features during model training, the attention analysis of **IDRBindNet** reveals that the model implicitly learns biologically meaningful signals associated with binding. Importantly, independent statistical analysis have been shown that sc_score is the single most strongly correlated descriptor with experimental *pK_d_*, and SHAP analysis of the Random Forest regressor likewise ranks sc_score and iupredB as the most influential predictors. The concordance between attention-derived correlations and these independent analyses provides strong evidence that the model learns representations aligned with intrinsic disorder and interfacial geometry in an unsupervised manner, indicating that these features play a dominant role in governing IDR-protein binding.

We have already demonstrated that model performance of **IDRBindNet** depends strongly on the choice of node embeddings, with one-hot encoding yielding the lowest performance and Prot-T5-BFD embeddings achieving the highest *R*^2^. To further disentangle the contribution of additional graph features, we investigated the role of edge features in determining model performance. The graph incorporates four edge features: C*α*-C*α* pairwise distance, orientation angle, difference in C*α* chemical shifts, and difference in SASA. To assess the relative importance of each of these features, we performed an ablation study in which each edge feature was individually removed by setting its values to zero, while keeping all other features unchanged. Model performance was then evaluated using PCC and *R*^2^ as done previously.

Overall, the ablation results (see Table S2) indicate that **IDRBindNet** is fairly robust to the removal of any single edge feature, suggesting a degree of redundancy among them. Nevertheless, removing the distance and chemical shift features leads to a noticeable drop in performance compared to the full model, indicating that these features contribute more substantially to predictive accuracy. In contrast, ablating the orientation angle has minimal impact on performance, implying that this feature provides comparatively less unique information in the presence of the other edge descriptors. The SASA difference shows an intermediate effect, contributing to performance but not as strongly as distance or chemical shift. Taken together, these results suggest that while the model leverages all edge features, geometric distance and chemical shift information play a more dominant role in capturing the interactions relevant for prediction, whereas orientation and SASA provide complementary role.

### Physicochemical Determinants of IDR-Protein Interactions

Given that shape complementarity between IDRs and their protein partners is a key determinant of binding affinity, we investigated several physicochemical properties (see “Methods”) to better understand the underlying forces governing these interactions. Our analysis revealed a strong negative correlation between the total solvent-accessible surface area of IDRs and shape complementarity (see Figure 5(a)). This indicates that the most successful IDR binders in our database are short, compact motifs. We think that smaller IDRs lose relatively little conformational entropy upon binding, allowing them to more readily achieve a precise interfacial fit with high shape complementarity and strong binding affinity. In contrast, larger IDRs possess greater inherent conformational entropy and therefore incur a substantial entropic penalty during association, as they must restrict many degrees of freedom to form a tight and well-defined interface.

**Figure 5:**
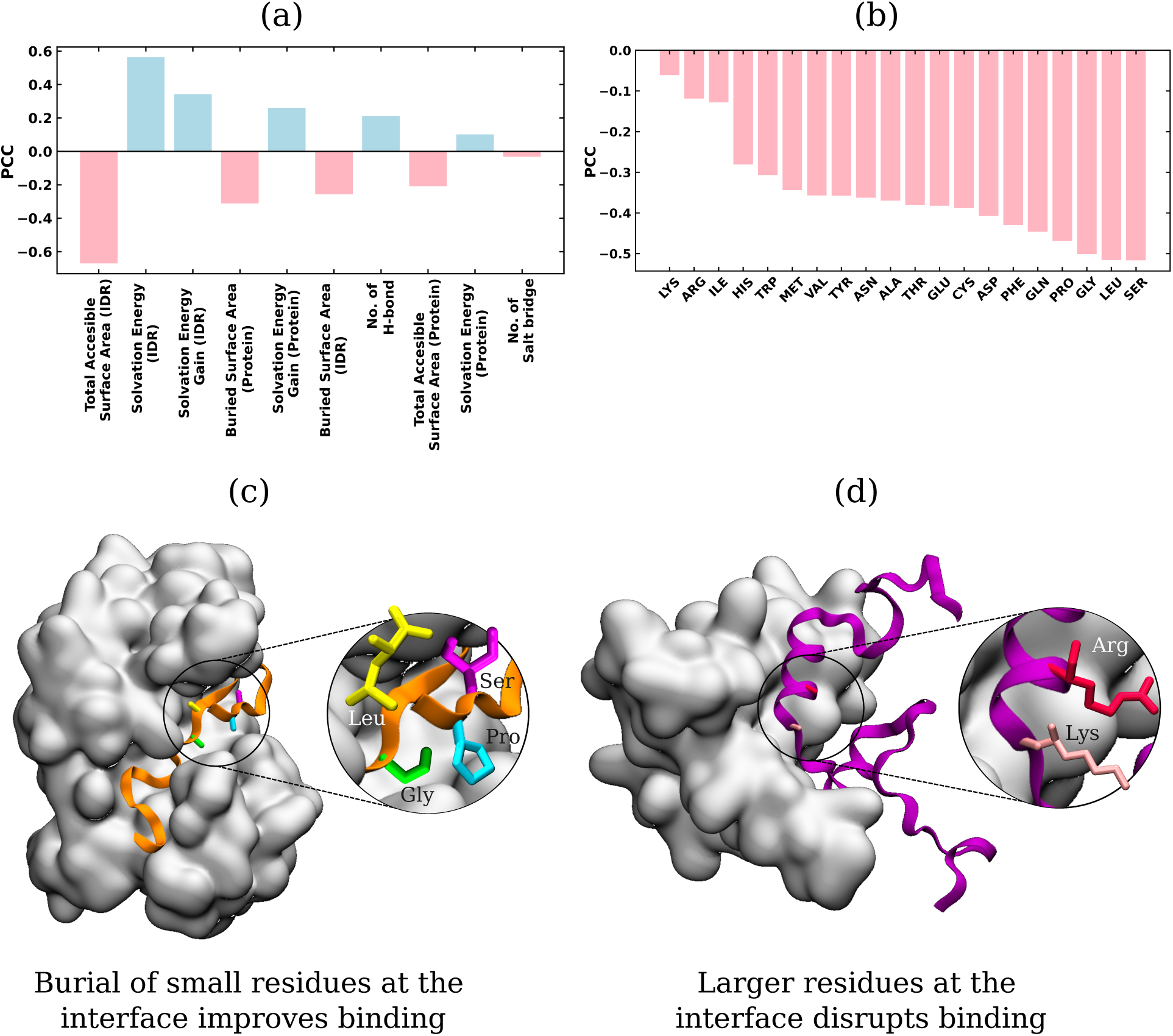
(a) Correlation of physicochemical properties of the interface with sc_score. (b) PCC of residuewise Total SASA with sc_score. (c) Schematic representation of a strong binding between IDR and protein and highlighting the key residues that contribute to high shape complementarity at the binding interface. (d) Schematic representation of a weak binding between IDR and protein and illustrating the residues that disrupt shape complementarity and reduce binding affinity.

A residue-level analysis of the binding interface reveals a clear hierarchy in amino acid contributions to shape complementarity. Burial of small and hydrophobic residues exhibits the strongest correlations with the sc_score, with serine, glycine, leucine, and proline emerging as the primary contributors (see Figure 5(b)). This observation is consistent with the recent findings reported by Balbi et al on surfaceomes, ^52^ who showed that leucine, valine, and serine are among the most frequently enriched residues in the predicted binding sites. In contrast, charged residues such as lysine and arginine show little to no correlation with sc_score (see Figure 5(b)). These results suggest that steric complementarity constitutes a fundamental constraint on interface formation in IDRs. We propose that enrichment of small residues particularly glycine and serine provides steric flexibility and spatial accommodation, enabling the IDR to approach the protein surface more closely. Such proximity would be sterically hindered by bulkier residues, thereby limiting optimal interfacial packing.

## Discussion

In this study, we constructed a comprehensive, high-quality dataset of 1,785 IDR-ordered protein interactions with experimentally determined *K_d_*, referred as *IBPC* − *K_d_*data-sets, addressing a critical gap in the availability of quantitative affinity data for disordered protein complexes. Using this dataset, we systematically analyzed the physicochemical and structural determinants of binding affinity, revealing that shape complementarity at the interface, structural order of the folded partner, and electrostatic complementarity are the dominant global features governing IDR-protein interactions.

We further developed a novel graph-transformer model, referred as **IDRBindNet**, that integrates protein language model embeddings with structural graph representations to predict *pK_d_*. The model achieved state-of-the-art predictive performance (PCC up to 0.956, *R*^2^ up to 0.911), significantly outperforming both traditional linear models and existing affinity prediction methods. Importantly, the model’s attention mechanism implicitly learned biologically meaningful properties, such as interfacial geometry and partner disorder, without explicit feature engineering, demonstrating its ability to capture the physical principles underlying binding. Finally, to make **IDRBindNet** broadly usable, we provide an easy-torun implementation on GitHub: users can obtain a predicted *K_d_*by simply supplying the sequences and structures of the two binding partners.

Unsupervised clustering of the feature space revealed distinct interaction regimes characterized by different combinations of physicochemical properties, which correlate with affinity ranges and reflect the diversity of IDR binding modes. Ablation studies confirmed the importance of geometric and chemical-shift edge features in the graph representation, while highlighting the complementary role of solvent accessibility and orientation information. Residue-level analysis of shape complementarity reveals that the burial of small, polar residues-particularly serine is highly sensitive to precise interfacial packing, underscoring its important role in modulating IDR protein interactions.

Together, this work provides a unified computational and biophysical framework for understanding and predicting IDR-protein binding affinities. The curated dataset, analytical insights, and predictive model establish a foundation for future studies rational design of IDR binders, and the exploration of disorder-mediated interactions in health and disease.

## Methods

### Construction of IDR-protein binding affinity dataset *IBPC* − *K_d_*

We started our investigation by basing on the DIBS^25^ database. While this database enlists 772 entries, only 487 of these entries have experimentally reported *K_d_* values. Starting from this database, we expanded the collection which includes specialized studies on model systems and are described below.

#### Baker *Nature* dataset^11^

Protein sequences were obtained from Supplementary Table-1 of the publication, and the corresponding affinity values were extracted from the main text. This dataset contains 11 IDR-protein pairs.

#### Baker *Science* dataset^10^

The *K_d_* values were collected from Supplementary Table-4 and Tables-S3A and S3B, while the IDR and binder protein sequences were extracted from the same study. This dataset comprises 39 samples.

#### Calcineurin dataset^26^

The IDR sequences and their corresponding *K_d_* values were obtained from Figure 5–figure supplements 1, 2, 5, and 6. The binder protein consists of calcineurin A (UniProt ID: Q08209, truncated to residues 1-400) and calcineurin B (UniProt ID: P63098). Several IDR sequences appeared multiple times in the supplementary data; for these duplicates, the reported *K_d_* values were averaged, while unique entries were retained as reported. Between calcineurin A and B, chain A was selected as the binding partner, as it exhibited a higher average residue-wise contact frequency than chain B. In total, this dataset contains 80 samples.

#### N2P2 dataset^29^

The binder protein is Na(+)/H(+) exchange regulatory cofactor NHE-RF2 (UniProt ID: Q15599, residues 142–254). The IDR sequences and corresponding *K_d_* values were extracted from Tables S2, S3, S4, and S5. IDR sequences listed in Tables S4 and S5 that were already present in Tables S2 and S3 were excluded to avoid redundancy. Specifically, the excluded sequences were AFDKYL, SFDKWL, and SFDKYL (from Table S4), and FFDTRF, FFDTRL, AFFTRL, FFSTRL, SFSTRF, ASFTRF, AFDKYL, FFDKYL, HSDFWL, SFDKYL, and SSFKYL (from Table S5). This dataset contains 47 samples.

#### DREB2A (Communication Biology) dataset^28^

The IDR consists of DREB2A and its mutants as reported in Table 1 (UniProt ID: O82132, residues 244–272). The binder protein is RCD1-RST (UniProt ID: Q8RY59, residues 499–572). This dataset includes 5 samples.

#### DREB2A (JACS) dataset^27^

Different fragments of DREB2A, as described in Table 3 of the publication, were used as IDRs. The binder protein RCD1-RST is identical to that used in the *Communication Biology* study. This dataset contains 4 samples.

#### STAMMPPING dataset^30^

The *K_d_*values were obtained from STAMMPPING experiments. The IDRs correspond to activation domains (ADs) derived from human transcription factors. A total of 374 ADs were previously identified, from which a subset of 204 was selected for STAMMPPING experiments. The binder proteins are human transcriptional co-activators; although 13 were initially tested, only 8 yielded high-confidence scores and were retained. The *K_d_* values were extracted from Supplementary Figure 2. The eight co-activators used are BRD7 (UniProt ID: QNPI1), P300 (UniProt ID: Q09472), P300KIX (residues 566–646), P300-TAZ1 (residues 332–417), P300-TAZ2 (residues 1726–1806), P300-NCBD (residues 2059–2117), TAF12 (UniProt ID: Q16514), and TAF6L (UniProt ID: Q9Y6J9).

For IDR sequence extraction, the DNA sequences provided for the activation domains were transcribed to obtain the corresponding protein sequences. Only ADs with unique DNA sequences were retained. All resulting IDRs are 80 residues long, consistent with the experimental report. The *K_d_* values were measured using either one or two experimental devices. When measured using a single device (P300, P300-KIX, P300-TAZ2, and TAF6L), the reported value was retained. When measured using two devices (BRD7, P300-NCBD, P300-TAZ1, and TAF12), the *K_d_* values were averaged. Extremely high reported values (e.g., ∼ 50,000 *µ*M) were discarded. The final number of samples for each co-activator is as follows: 167 (BRD7), 157 (P300), 108 (P300-KIX), 135 (P300-NCBD), 137 (P300-TAZ1), 120 (P300-TAZ2), 165 (TAF12), and 123 (TAF6L).

In total, the curated dataset *IBPC* − *K_d_* comprises 1,785 IDR-protein interaction pairs with experimentally determined *K_d_* values and is represented in the pie chart as shown in Figure 1 (a). The corresponding IDR-protein sequence pairs were used as input to AlphaFold 3^53^ for structure prediction. For each interaction, the predicted complex structure with the highest ranking score was selected for all subsequent analyses. The *K_d_* distributions are visualized using kernel density estimation (KDE) performed in logarithmic space to account for the wide dynamic range of binding affinities. For each dataset, the *K_d_* values are transformed as log_10_(*K_d_*). A Gaussian KDE is then fitted to the log-transformed values and evaluated on a dense grid spanning affinities from 10 mM to 1 pM. The resulting density is plotted against the corresponding linear-scale *K_d_* values, producing a smooth probability density curve that reflects the relative likelihood of observing a given binding affinity in log space with weaker binding affinities (higher *K_d_*) appear on the left and stronger affinities (lower *K_d_*) appear on the right (see Figure 1 (b)).

### Unsupervised Clustering of IDR-protein features

The 15-dimensional feature space was reduced to two dimensions using Principal Component Analysis (PCA), t-distributed Stochastic Neighbor Embedding (t-SNE), and Uniform Manifold Approximation and Projection (UMAP) (see Figure 2(a)). Each embedding was independently clustered using Gaussian Mixture Models (GMMs) across a range of cluster numbers (*k* = 2 − 20), with five random initializations per cluster. Among the three methods, UMAP consistently produced higher silhouette scores than t-SNE and PCA, indicating superior separation of latent structure (see Figure 2 (b)). For the UMAP embedding, silhouette scores exhibited pronounced local maxima at *k* = 4 and *k* = 9, while the Bayesian Information Criterion (BIC) displayed an inflection in its gradient near *k* = 8, beyond which additional clusters yielded diminishing improvements in model fit. Considering both metrics, we selected *k* = 9 as the optimal partitioning, which was adopted for all subsequent analyses.

### Supervised regression models for binding affinity prediction

To evaluate the predictive power of the 15 engineered physicochemical and structural descriptors, we trained several supervised regression models to predict binding affinity (*pK_d_*). The *pK_d_* values were scaled down further by the total number of residues in the complex. All input features and the target variable were standardized to zero mean and an unit variance. The dataset was split into training (80%) and test (20%) sets with a fixed random seed. We compared linear models (Linear Regression baseline, Lasso, ^45^ and Ridge regression^46^ with *α* tuned via 5-fold cross-validation over logarithmically spaced grids: 10*^−^*^4^-10^1^ for Lasso and 10*^−^*^4^-10^3^ for Ridge) against nonlinear ensemble models (XGBoost ^47^ and Random Forest^48^), with hyperparameters optimized via grid search and 5-fold cross-validation. Feature importance was interpreted using SHAP^49^ (SHapley Additive exPlanations) values, with mean absolute SHAP values used to rank global feature importance.

### Graph representation of IDR-Protein complex

Node features are derived from contextual embeddings obtained from multiple pretrained protein language models (pLMs). The primary amino acid sequences of the IDR and the binder protein in *IBPC* − *K_d_*database are first concatenated into a single continuous sequence, which is then fed into several pLMs, including Evolutionary Scale Modeling (ESM-2)^50^ variants with 8M, 35M, 150M, 650M, and 3B parameters, ProtT5, ^51^ ProstT5,^54^ ProtBERT,^51^ and ProtT5-BFD.^51^ The resulting per-residue embeddings are subsequently separated again into distinct sets corresponding to the residues of the IDR and the binder protein, respectively.

We used AlphaFold3^53^ to predict the structure of each IDR-binder complex. For the residue-level graph, we extracted four structurally and biophysically informative edge features directly from the AlphaFold3-predicted complexes.: (i) the pairwise C*α*–C*α* distance, (ii) the relative spatial orientation between residues, (iii) the difference in C*α* chemical shifts, and (iv) the residue-wise difference in solvent-accessible surface area (SASA). Edges were defined between residue pairs based on distance thresholds applied to the C*α*-C*α* separation. A cutoff of 6 Å was used for intramolecular residue pairs, while a cutoff of 8 Å was applied to intermolecular residue pairs between the IDR and its binding partner. MDTraj^55^ was employed for extracting features (i) and (iv). The details of other feature extraction process is provided below:

To encode the relative spatial orientation between residues in a graph representation of the complex structure, we derive rotation-invariant edge features based on the local coordinate frames of backbone atoms. Each residue is represented by its three backbone atoms: the amide nitrogen (**N**), alpha carbon (**C***α*), and carbonyl carbon (**C**). For each residue *i*, a local orthonormal coordinate frame is constructed using Gram-Schmidt orthonormalization. Given the 3D coordinates of the backbone atoms **N***_i_*, **C***α_i_*, and **C***_i_*, two vectors are defined as follows:

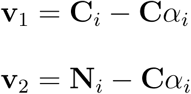

The first basis vector is obtained by normalization:

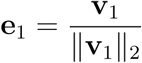

The second vector is orthogonalized against **e**_1_:

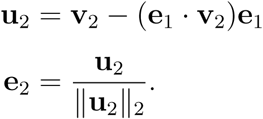

The third basis vector is obtained via the cross product to complete the right-handed orthonormal frame:

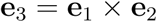

The resulting rotation matrix **R***_i_* ∈ SO(3) for residue *i* is formed by stacking the basis vectors as columns:

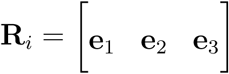

This procedure is applied independently to every residue in the IDR and the protein from the AF3 structure. For each complex model, all pairwise relative rotation matrices **R***_kl_* between residues *k* and *l* are computed as

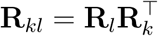

This quantity is invariant to global rotation of the entire structure and captures the relative orientation of the local peptide planes. To convert these relative rotations into a scalar edge feature suitable for graph neural networks, the rotation angle *θ_kl_* ∈ [0*, π*] is extracted from each **R***_kl_* using the trace formula for rotation matrices:

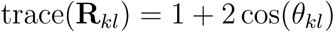

Solving for the angle yields

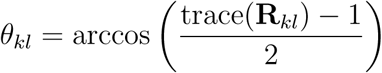

This procedure is repeated across all generated structural models, yielding one angle matrix per complex. The computed angle features are incorporated as edge attributes in the graph representation. This was derived by in-house Python code developed by the authors.

Chemical shifts are highly sensitive to the local chemical environment, secondary structure, and conformational state of each residue. For each residue pair *k* and *l* connected by an edge, the scalar edge feature is computed as the absolute difference in their C*α* chemical shifts (obtained from SPARTA+^56^)

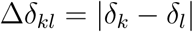

where *δ_k_* and *δ_l_* denote the C*α* chemical shift (in ppm) for residues *k* and *l*, respectively. This difference encodes local environmental dissimilarity and potential conformational perturbations induced by binding or structural context, providing complementary physicochemical information to purely geometric features.

The per-residue SASA quantifies the extent of solvent exposure and is typically reduced for residues participating in binding interfaces. We compute per-residue SASA values from the AF3-predicted complex structures using the Shrake-Rupley algorithm as implemented in MDTraj,^55^ with a probe radius of 1.4 Å. The SASA-based edge feature between residues *k* and *l* is defined as the absolute difference

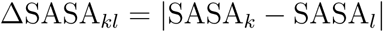

This pairwise measure captures relative differences in residue burial and solvent exposure, thereby emphasizing potential interaction hotspots where one residue becomes significantly more buried than its neighboring partner across the interface. As a result, this feature is particularly effective in distinguishing intrachain packing interactions from intermolecular contacts in IDR-protein complexes. This provides the basic constituent of a graph transformer model (hereby coined as **IDRBindNet**), as described in following paragraph.

### Model architecture and training protocol of IDRBindNet

**IDRBindNet**, the Graph Transformer model for Regression processes molecular graphs, where each node is initialized with a *d*_LLM_-dimensional embedding obtained from a pretrained large language model, and each edge is described by a 4-dimensional feature vector. The raw edge features are first projected into the model’s hidden dimension *d*_h_ = 128 through a two-layer MLP consisting of a linear layer reducing to 64 units, followed by ReLU^57^ activation and a final linear projection to *d*_h_. Node representations are then updated by three successive TransformerConv^58^ layers that incorporate these projected edge embeddings in the attention mechanism.^59^ The first two layers each employ 8 attention heads with concatenation (producing an intermediate representation of size 8 × 128), while the third layer uses a single attention head without concatenation (output size 128). Each TransformerConv layer is followed by batch normalization matched to its output channel size, an ELU^60^ activation function, and dropout^61^ with probability 0.1. After the final transformer layer, node embeddings are aggregated into a graph-level representation using global mean pooling over all nodes in the graph. The resulting fixed-size graph embedding (128-dimensional) is passed through a three-layer feed-forward regression head consisting of successive linear transformations 128 → 64 → 16 → 1, with ReLU activations applied after the first two linear layers, yielding the final scalar prediction.

For graph construction, both node embeddings and edge features were standardized. For prediction, the *pK_d_* values were rescaled by the total number of residues in the IDR-protein complex. The *IBPC* − *K_d_*dataset was split into training, validation, and test sets in a 70:15:15 ratio, ensuring that each split contained at least one sample from each source from which this dataset was curated. Five independent dataset splits were generated. The model was trained for 300 epochs using a batch size of 32 and the Adagrad^62^ optimizer with a learning rate of 10*^−^*^2^. The mean squared error (MSE) was used as the loss function. Model performance was evaluated on the test set using the Pearson correlation coefficient (PCC) and the coefficient of determination (*R*^2^), reported as the mean and standard deviation across the five dataset splits. The model, referred as **IDRBindNet**, is publicly available online (https://github.com/JMLab-tifrh/IDRBindNet). Given the structural coordinate of the IDR-protein complexes, this model will provide an estimate of its *K_d_* in few minutes.

### Dataset Splitting and Construction of the out-of-distribution (OOD) Benchmark

To evaluate model generalization on sequence dissimilarity, we implemented a clustering-based out-of-distribution (OOD) data split strategy. Protein sequences were first clustered using CD-HIT^63^ with a sequence identity threshold of 40%, requiring a word size of 2 for matching. This threshold ensures that sequences within the same cluster share ≥40% sequence identity, while sequences from different clusters share *<*40% identity and are therefore considered evolutionarily distant. Following clustering, each sequence was assigned to a cluster based on the CD-HIT output. To create OOD train/test splits, we employed grouped cross-validation using GroupKFold (5-fold) from scikit-learn, ^64^ with cluster assignments as the grouping variable. This ensured that all sequences from the same cluster were placed exclusively in either the training or test set within a fold, preventing data leakage and simulating a realistic scenario in which test sequences have no close homologs (≥40% identity) in the training data. For each fold, we verified zero cluster overlap between training and test sets and the resulting datasets were used for subsequent model training and evaluation.

### Extraction of Interfacial Physicochemical Features

To quantitatively characterize the physicochemical properties at IDR-protein interfaces, we used PISA v.2.2.0 (Protein Interfaces, Surfaces and Assemblies) within the CCP4 suite^65^ to extract a comprehensive set of features for each complex in our database. For every structure, we obtained the total accessible surface area, solvation energy, solvation energy gain upon binding, and buried surface area for both the IDR and its protein partner. Additionally, we extracted the number of intermolecular hydrogen bonds and salt bridges formed at the interface.

## Code and Data Availability

The code of our model (**IDRBindNet**) for predicting binding affinity can be accessed on GitHub via the following link: https://github.com/JMLab-tifrh/IDRBindNet. The **IBPC-***K_d_* dataset curated in this project, comprising of sequences and structures of IDR and its partner protein along with experimentally measured *K_d_* has been uploaded in Zenodo and can be accessed via the following link: https://zenodo.org/records/18722319.

## Supporting Information

The Supporting Information (SI) provides supplemental figures, supplemental table. Figure S1 shows the correlation between the features and *pK_d_* values for DIBS and Baker’s data. Figure S2 shows the latent structure of the dataset using PCA and t-SNE. Figure S3 shows the correlation of z-score of different features with mean *pK_d_*. Figure S4 shows the *R*^2^ value for Linear, Lasso, Ridge, XGB and RF regression models. Figure S5 shows the AF3 predicted complex from external dataset along with the experimental and predicted *K_d_*. Table S1 shows the PCC and *R*^2^ for different PLM’s and other *K_d_* prediction models. Table S2 presents the ablation study of edge features for the best-performing model. Table S3 shows the comparison between predicted and experimental *K_d_* values from an external source.^52^

## Author Contributions

SA curated the dataset, built the graph-transformer model and performed the training and evaluations. SC performed the benchmarking with other ML models. SA and JM surveyed literature, analysed the data and co-wrote original and revised manuscript. JM supervised the investigation and acquired the funding.

## Acknowledgement

We acknowledge support of the Department of Atomic Energy, Government of India, under Project Identification No. RTI 4007. We sincerely acknowledge Tata Institute of Fundamental Research Hyderabad, India for providing the support of computing resources. JM acknowledges the core research grant approved by the Department of Science and Technology (DST) of India (CRG/2023/001426).

## Notes

### Competing Interest Statement

The authors have declared no competing interest.

