## Supplemental figures and tables for "A Comprehensive Atlas and Machine-Learning Framework for Predicting IDR-Protein Binding Affinity"

*36/P, Gopanpally Village, Serilingampally Mandal, Ranga Reddy District, Hyderabad 500046, India*

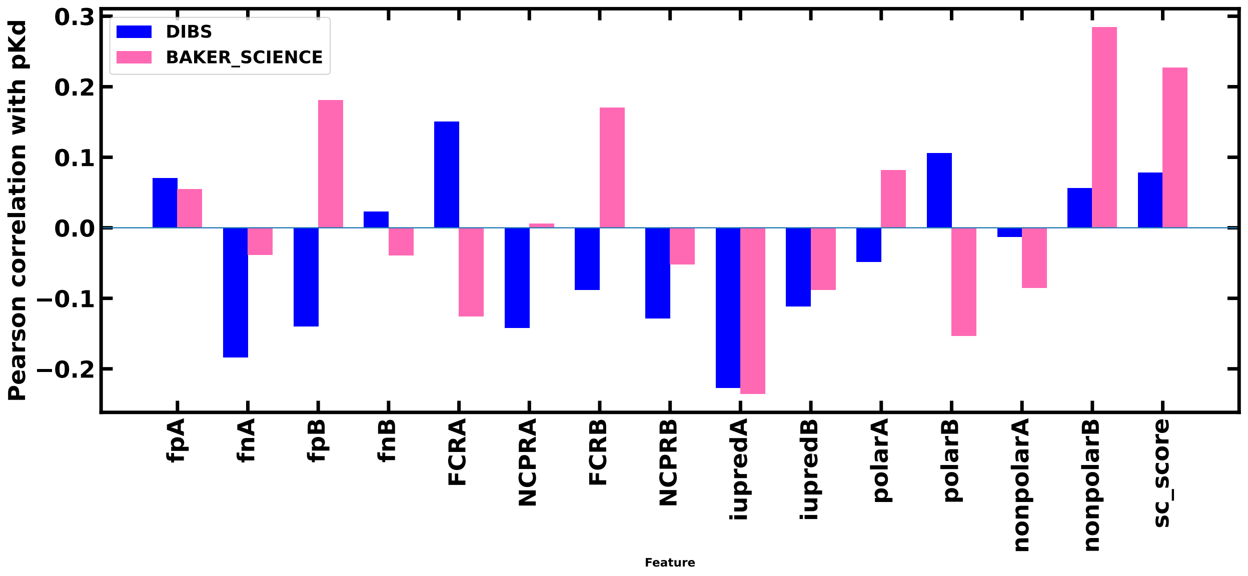


**Figure S1:** Correlation of the features with $pK_d$ for DIBS data (reference 25) and the Baker (*Science*) (reference 10) data

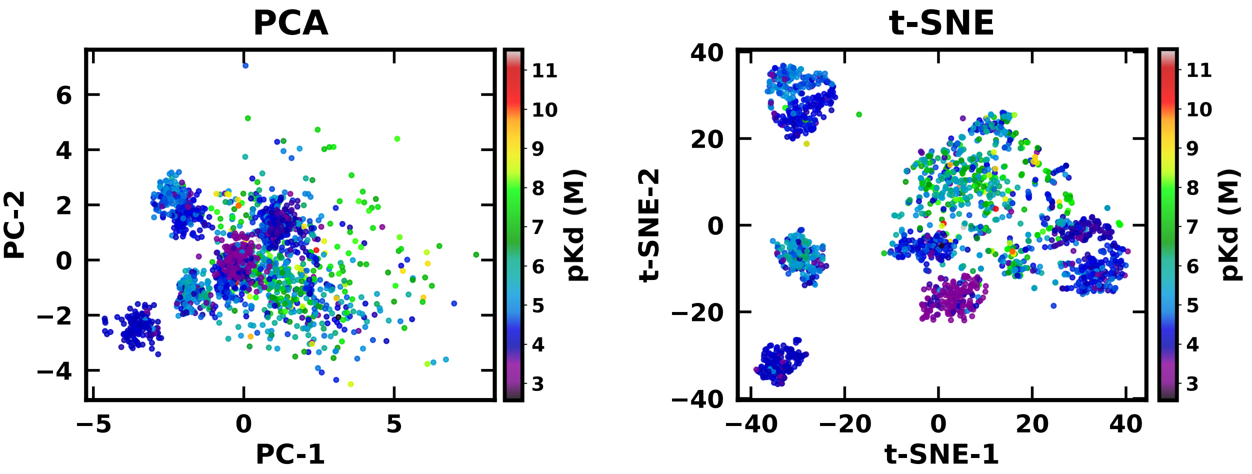


**Figure S2:** Latent structure of the dataset visualized using (a) PCA, (b) t-SNE

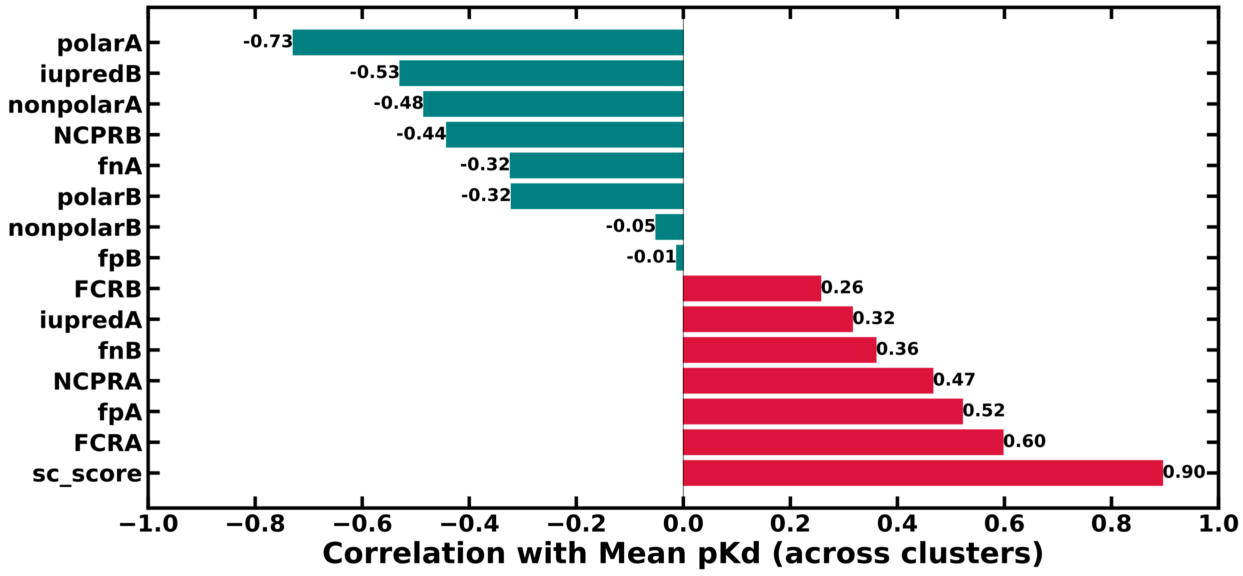


**Figure S3:** Correlation of z-score of different features with mean $pK_d$.

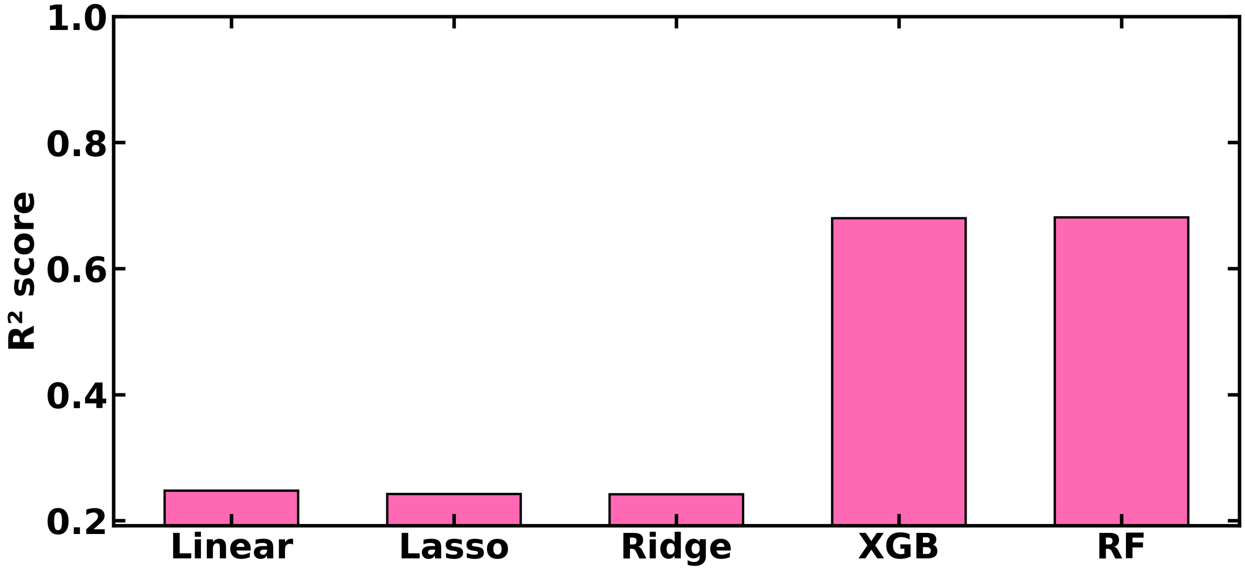


**Figure S4:** Comparison of $R^2$ values for regression models predicting $pK_d$

**Table S1:** Benchmarking of our model with reported model on our internal test set

| Model | PCC | $R^2$ |
|---|---|---|
| ONE HOT | $0.924 \pm 0.011$ | $0.850 \pm 0.020$ |
| ESM2-8M | $0.929 \pm 0.013$ | $0.859 \pm 0.024$ |
| ESM2-35M | $0.931 \pm 0.011$ | $0.865 \pm 0.020$ |
| ESM2-150M | $0.935 \pm 0.015$ | $0.860 \pm 0.040$ |
| ESM2-650M | $0.939 \pm 0.013$ | $0.880 \pm 0.025$ |
| ESM2-3B | $0.947 \pm 0.009$ | $0.895 \pm 0.018$ |
| PROST-T5 | $0.953 \pm 0.007$ | $0.906 \pm 0.013$ |
| PROT-T5 | $0.954 \pm 0.007$ | $0.909 \pm 0.014$ |
| PROT-BERT | $0.927 \pm 0.019$ | $0.853 \pm 0.034$ |
| **PROT-T5-BFD** | $\mathbf{0.956 \pm 0.003}$ | $\mathbf{0.911 \pm 0.007}$ |
| PPAP | $0.924 \pm 0.011$ | $0.582 \pm 0.041$ |
| ProAffinity-GNN | $0.888 \pm 0.014$ | $0.697 \pm 0.036$ |

**Table S2:** Ablation study

| Model | PCC | $R^2$ |
|---|---|---|
| Distance | $0.954 \pm 0.004$ | $0.898 \pm 0.008$ |
| Orientation angle | $0.958 \pm 0.006$ | $0.913 \pm 0.011$ |
| Chemical Shift | $0.957 \pm 0.006$ | $0.902 \pm 0.013$ |
| SASA | $0.957 \pm 0.006$ | $0.906 \pm 0.015$ |

**Table S3:** Experimental and predicted $K_d$ values from external dataset (see reference 52). The predicted values are from the best model.

| Complex | Experimental | Predicted |
|---|---|---|
| F_rd1_702 | 741 nM | 260 nM |
| F_rd2_046 | 147 nM | 41 nM |
| I_rd2_159 | 2.5 $\mu$M | 7.3 $\mu$M |
| F_rd2_037 | 172 nM | 142 nM |
| H_rd1_309 | 34 $\mu$M | 407 $\mu$M |

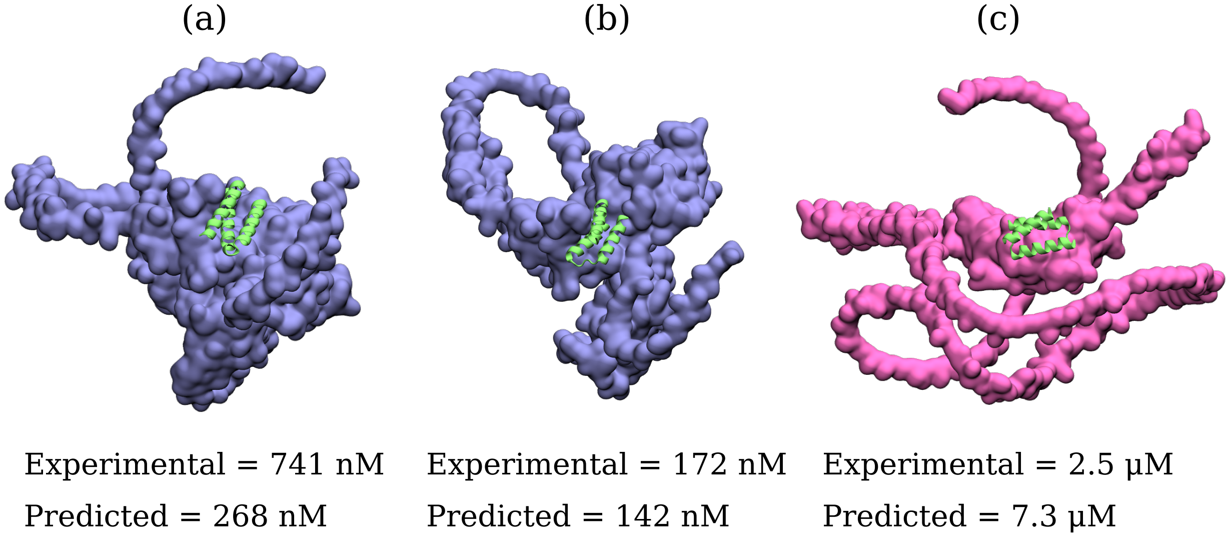


**Figure S5:** (a)-(c) External validation on *de novo* designed binders of a set of surface-bound proteins: AlphaFold3-predicted complex structures for F_rd1_702, (d) F_rd2_037 and (e) I_rd2_159 reported by Balbi *et al.*, shown alongside the experimentally measured dissociation constants ($K_d$) and the corresponding $K_d$ values predicted by our model. See Table S3 for comparison on a set of five designed protein complexes from the report by Balbi *et al.* (Reference 52)
